# *Chlamydia trachomatis* genomes from rectal samples: description of new clade comprising *ompA*-genotype L4 from Argentina

**DOI:** 10.1101/2024.11.23.624510

**Authors:** Karina Andrea Büttner, Fanny Wegner, Vera Bregy, Andrea Carolina Entrocassi, María Lucía Gallo Vaulet, Deysi López Aquino, Luciana La Rosa, Laura Svidler López, Mirja H Puolakkainen, Eija Hiltunen-Back, Frank Imkamp, Adrian Egli, Helena MB Seth-Smith, Marcelo Rodríguez Fermepin, ESGMAC

## Abstract

Whole genome analysis has provided us with insights into the evolution of *Chlamydia trachomatis* and recently into circulating strains which cause lymphogranuloma venereum (LGV). A large LGV outbreak of a new *ompA*-genotype, L2b, was first reported in Europe in the early 2000s, primarily affecting men who have sex with men (MSM), and then expanded globally. More recent work shows this outbreak diversifying into variants of described *ompA*-genotypes, with the same L2b genomic backbone. This study extends the investigation of LGV cases to Argentina and Finland. In 2017, an LGV outbreak was described in Argentina characterized by distinct genomic features shown by both *ompA*-genotyping and MLST analysis. We have obtained whole genome sequences from cultured isolates and clinical samples via SureSelect (Agilent) target enrichment. Based on *ompA* and phylogenetic analyses, we describe further diversity within the *ompA*-genotype L2b clade, illustrating the transmission dynamics in both Argentina and Finland. A key finding is that of a novel clade of Argentinian samples, characterised by a proposed new *ompA*-genotype L4. Additionally, we present the genome sequence of a non-LGV strain associated with anorectal proctitis. These findings contribute to the investigation of LGV evolution, particularly with the presence of the novel L4 lineage, and provide insights into genomic diversity and transmission dynamics of *C. trachomatis*.

**Graphical Abstract:** 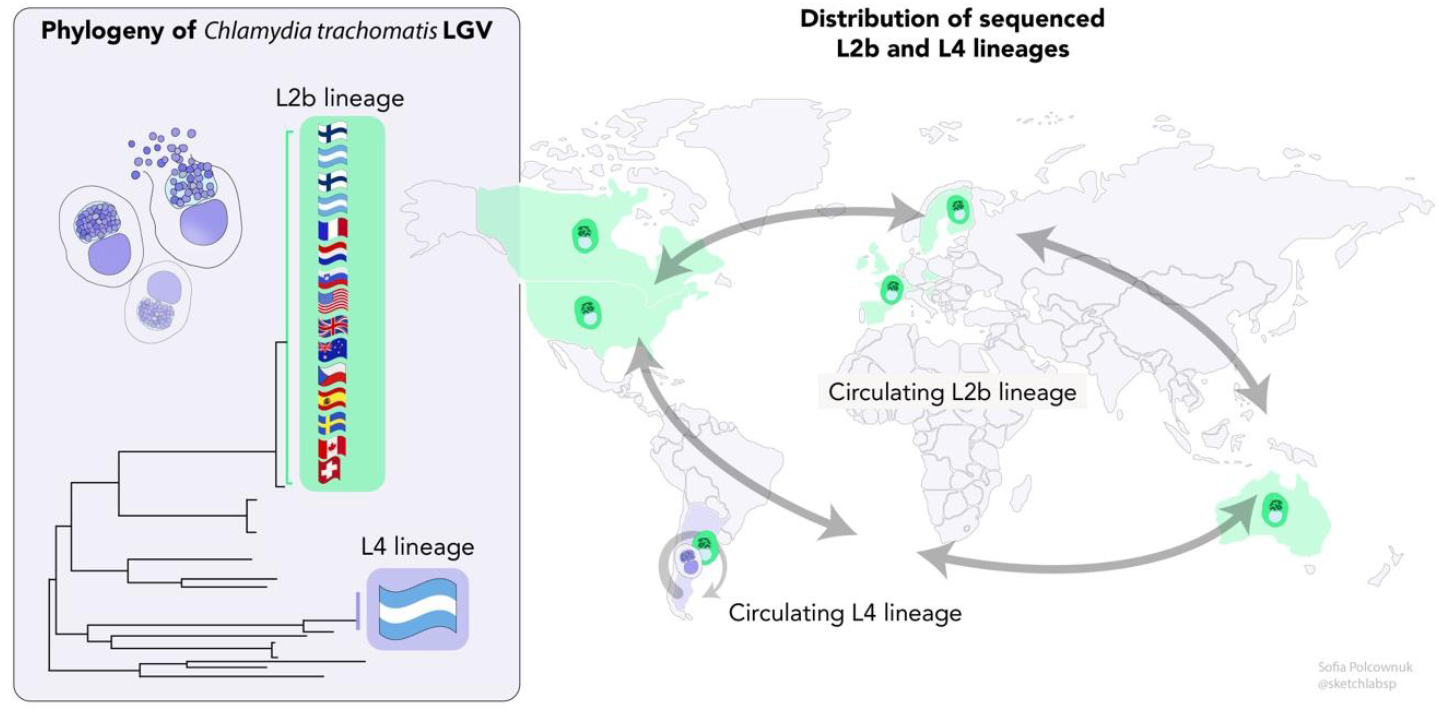

**Impact statement:** The phylogeny of *C. trachomatis* has been described in several publications and has been clear for a decade. The evolution of this STI in the intervening years also informs a great deal about the transmission opportunities and selective pressures that the intracellular bacterium is under. Through genome sequencing, also directly from clinical samples, we provide the first LGV genomes from Argentina and Finland, and describe further evidence of global circulation of the *ompA*-genotype L2b lineage. We describe a fully novel lineage of LGV *C. trachomatis*, from Argentina, proposed as *ompA*-genotype L4. We also find evidence of a strain causing proctitis from the urogenital lineage. Together, these provide significant new findings in the investigation of *C. trachomatis*.

**Data summary:** All illumina sequence data, with human read data removed using Hostile (1) and KrakenTools (https://github.com/jenniferlu717/KrakenTools), is deposited with the European Nucleotide Archive (ENA) under project number PRJEB72167.

## Introduction

The sexually transmitted infection (STI) causing bacterium, *Chlamydia trachomatis* has an invasive biovar, LGV, causing lymphogranuloma venereum, as well as the urogenital (UG) and ocular biovars. Recently, interest has arisen around the current evolution of LGV strains of *C. trachomatis* (2–4). To further investigate global circulating strains of *C. trachomatis* LGV, whole genome sequencing is the best method.

The most recent genomic findings on LGV *C. trachomatis* showed ongoing evolution of the *ompA*-L2b lineage, with the few genomic single nucleotide polymorphisms (SNPs) found over-represented in the *ompA* gene, generating novel variations in the encoded MOMP protein (2).

Comprehensive nationwide prevalence data for *C. trachomatis* is lacking in Argentina, as is identification of all LGV cases, although the limited data suggest higher rates of LGV in Buenos Aires than in Europe (5–7). Since 2017, proctitis patients in Buenos Aires, mostly HIV-infected MSM, have been screened for *C. trachomatis* (8), showing a range of *ompA*-genotypes (9) and MLST types (10).

This work further investigates LGV genomes from Argentina and Finland in the context of global data, to broaden the spectrum of LGV genomes analysed to date. We show additional and further development in the *ompA*-genotype L2b lineage, confirm the genomic separation of a new South American lineage, termed *ompA*-genotype L4, and the context of a rectally derived UG lineage strain.

## Methods

### Sample collection, culture and DNA extraction

Rectal swab samples were obtained from symptomatic adult patients presenting proctitis-compatible symptoms and STI risk factors between September 2017 and August 2023 attending in a public hospital and in a private clinic in Buenos Aires, Argentina (Hospital Juan A. Fernández and Centro Privado de Cirugía y Coloproctología). Additionally, rectal swab samples from symptomatic patients and those with partner notifications attending the outpatient STI clinic at the Helsinki University Central Hospital (HUCH) in Finland between 2011-2013 were collected from MSM with symptoms suggestive of LGV or due to partner notification. These samples were either cultured and/or had DNA extracted as described in (11).

### Sequencing

Genomic sequencing data was obtained either from cultured samples or via SureSelect target enrichment (Agilent) (Table S1). Methods were performed as described in (11).

### Sequence analysis

Reads were trimmed with trimmomatic v0.39 (12). Mapping to reference AM884177, base calling and generating of multiple sequence alignment (MSA) used snippy v4.6.0 (https://github.com/tseemann/snippy) with default parameters except a minimum coverage of 5 to enable calling from regions of lower read depth, and with the rRNA operons masked. Genomes were considered acceptably complete when percentage genome covered >95%, mean read depth >10, and <20% Ns called at <0-5x read depth. Presence of more than one strain of *C. trachomatis* in all samples was analysed using the R package vcfR (13) by looking at all minority variants with a minimum frequency of 10% and a minimum depth of 5. All were found to have fewer than 200 minority variants and are thus considered to be pure strains.

Phylogenies were created using all LGV genome data available (2,14,15) in IQtree v2.2.0.3 (16) with 1000 bootstraps. Gubbins v3.3.0 (17) was used to remove recombinations, with 5 iterations, 1000 bootstraps, iqtree-fast as first-tree-builder, iqtree as tree-builder, and custom model K3Pu+F+I+I+R3. FastBAPS was used to define clades (https://github.com/gtonkinhill/fastbaps). UG phylogenies were built by comparing to previous UG genomes and mapping against the genome of E/Bour (HE601870) (14). Phylogenies were visualised with associated metadata in Phandango (18). The whole genome alignments are given in Suppl File1 and 2.

Complete CDS sequences of *ompA* were extracted from the mapped alignments, compared with capillary sequence data (10) and blastn results from the NCBI database in AliView v1.27 (19) (Table S1).

### Statistical analysis

Statistical analysis of patient data was performed using InfoStat software version 2020, Universidad Nacional de Córdoba, Córdoba, Argentina. The two-tailed Fisher exact test was used for categorical variable analysis. Statistical significance was defined as *p*<0.05.

## Results and Discussion

### Phylogeny of *C. trachomatis* LGV genomes

*C. trachomatis* LGV genomes (n=32 from Argentina and n=9 from Finland) were obtained following either culture (n=10 from Argentina and n=9 from Finland) or SureSelect (n=22 from Argentina, one of which had been cultured but required target enrichment to produce a full genome) and Illumina sequencing. These are the first genomes from Argentinian and Finnish LGV strains. The genomes were compared phylogenetically to previously published LGV genomes (Figure 1). A phylogeny with duplicate samples (n=8), showing the reproducibility of the SureSelect method is given in Figure S1, and the recombinations identified within the LGV genomes shown in Figure S2.

**Figure 1.**
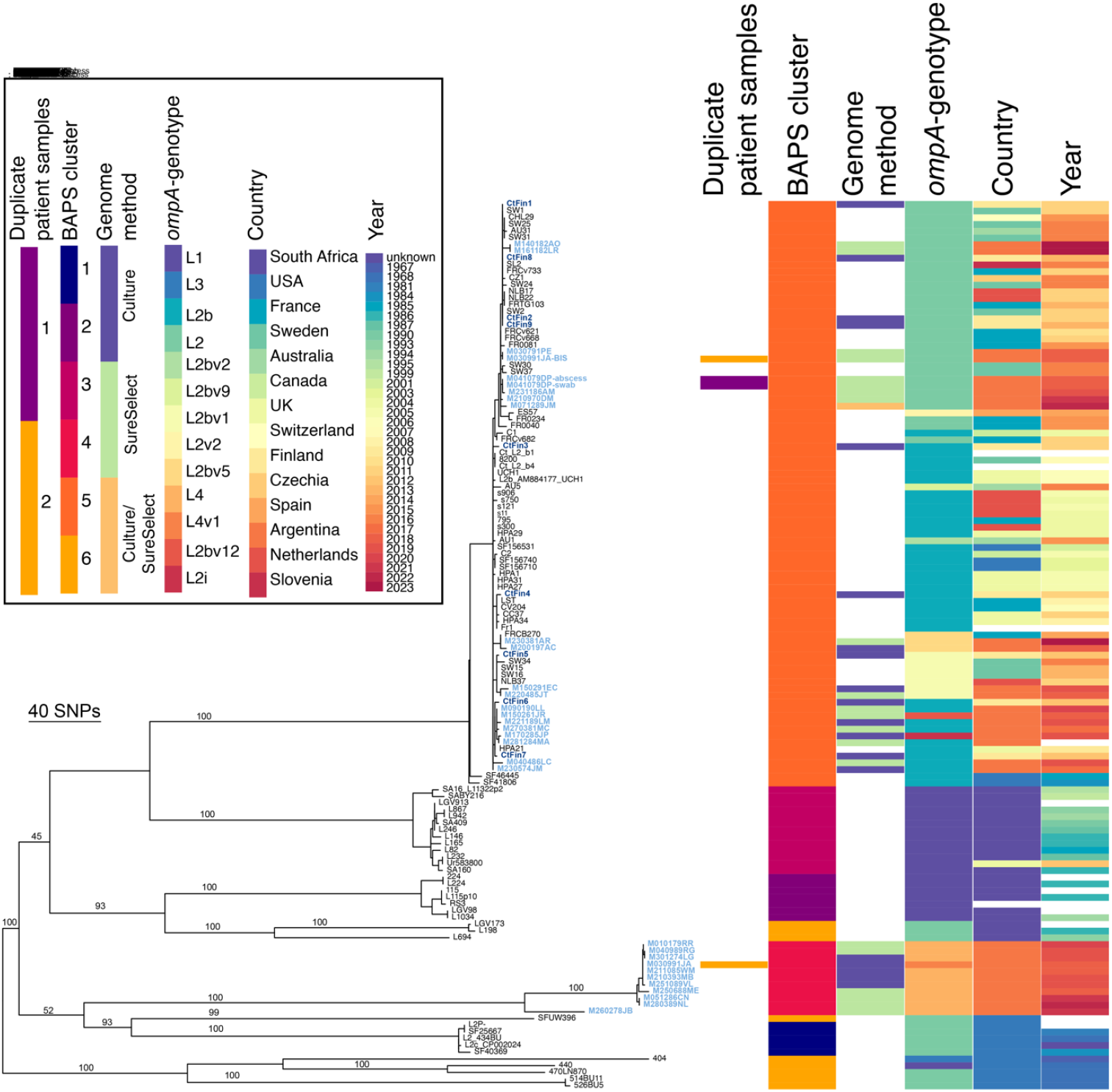
Recombination-adjusted phylogeny of LGV strains from Argentina, Finland and from previous publications. All published LGV genomes with available data were included in the analysis. Right of the phylogeny is relevant metadata: duplicate patient samples, BAPS cluster (clade), genome method (data for this paper), *ompA*-genotype, country, year as described in the key to the top left. Missing data is displayed as white. Names of genomes from this publication are coloured in the phylogeny in light blue from Argentina and in dark blue from Finland. Bootstrap values from 1000 bootstraps are shown as % shown on major branches.

### Further global circulation of the *ompA*-genotype L2b lineage

Many (n=21/32) of the Argentinian samples, and all the Finnish samples (n=9), fall within the *ompA*-genotype L2b clade (BAPS cluster 5 in Figure 1). The samples within this L2b clade show diverse *ompA*-genotypes even over short timespans, suggesting selective pressure on the encoded MOMP protein. Several sub-clades are apparent, with six separate locations of the Finnish samples in the phylogeny speaking for European circulation of many strains. Three pairs of samples represent possible local circulation of strains in Helsinki: these are unlikely to be from direct partners or through partner notifications, due to disparate sampling dates.

The closely related groups of two to six genomes from Argentinian samples suggest locally circulating strains among the MSM community in Buenos Aires. One patient had two genomically identical samples from two different locations (abscess and anorectal swab), both being *ompA*-genotype L2 within the L2b clade. It is worth noting that the other samples in sub-clades were independent, and based on the patients’ histories were likely not connected. These groups are often closely related to samples from Europe, indicating multiple importation of strains from diverse international sources. As these are the most recent samples in the phylogeny, ongoing travel is not yet apparent. Of note, two Argentinian samples from 2017 and 2023 respectively, carry identical genomes to several European samples including L2b/UCH1 and L2b/UCH2 from 2006, reiterating that *C. trachomatis* has a less clock-like mutation rate than other bacterial species (14). The L2b clade is clearly highly international, reflecting a global circulating lineage of *C. trachomatis* LGV.

Complete genomes were obtained of two samples carrying new *ompA* variants (L2i and L2bv12) recently described in Argentina (10). The *ompA* variant L2i, carrying a mutation which leads to an S162I substitution at the site that differentiates *ompA* genotypes L2 and L2b, and L2bv12, carrying a mutation that leads to the S164D substitution, both in the variable domain (VD)2.

### Description of novel lineage from Argentina

We identified a new lineage comprising 11 Argentinian samples. This separate clade (BAPS cluster 4 in Figure 1) is defined by a branch of >200 SNPs and is confirmed as a separate genetic cluster. Of interest, one patient had two samples taken over time: *ompA*-genotype L4 from April 2018, and *ompA*-genotype L2 within the L2b clade from August 2018, suggesting reinfection with a different strain after treatment (Figure 1).

The *ompA* gene of the L4 clade is most similar to that of *ompA*-genotype L1 (accession HE601950.1), but with ten nucleotide substitutions; two versions of *ompA*-genotype L4 (L4 and L4v1) vary by a single SNP (10) (Table S1). The amino acid substitutions compared to *ompA*-genotype L1 are all located in VDs: A90T in VD1, N162S in VD2, and T311A in VD4. Taking into account this level of diversity from all previously described LGV genomes, and the number of samples defining the clade (n=11), we propose the new *ompA*-genotype L4.

This novel lineage shows a tight sub-clade of strains presumably representing an ongoing outbreak, with a more distantly related member also represented towards the root of the clade, showing the diversity present in the Buenos Aires area. There is likely to be further unsampled diversity within this clade, perhaps with more diverse members of this clade in other areas of South America. The lineage also appears to be present in France (20): genomics could here give insights into the origins of this lineage.

**Figure S1.**
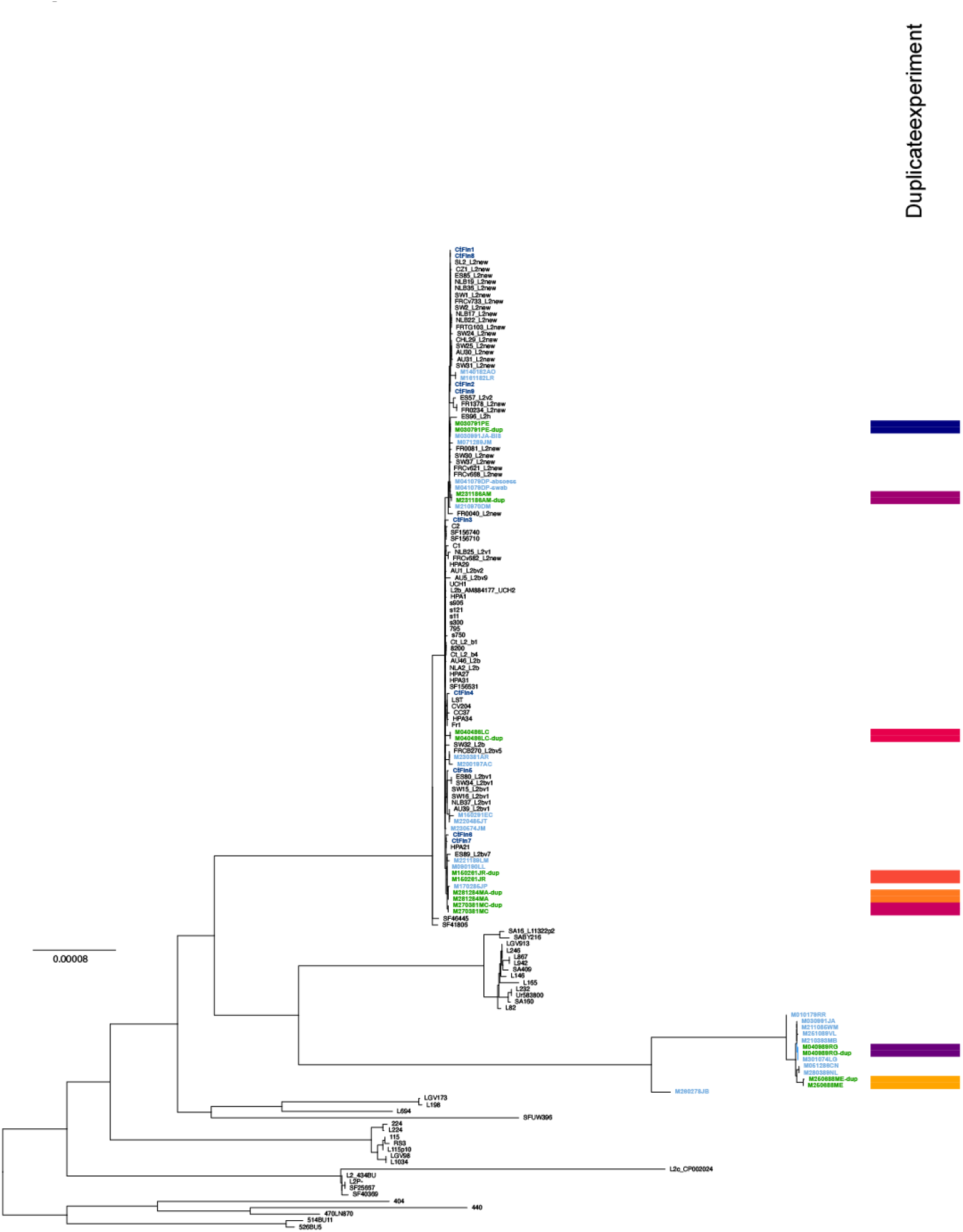
Phylogeny of LGV strains without adjusting for recombination, including duplicate experiments. Duplicate experiments (n=8, to the right of the phylogeny, paired colours) show the robustness of the SureSelect method. Names of genomes from this publication are coloured in the phylogeny in light blue from Argentina and in dark blue from Finland. Names in green are those with identical duplicates (“-dup”). Scale bar refers to a phylogenetic distance of 0.00008 nucleotide substitutions per site.

**Figure S2.**
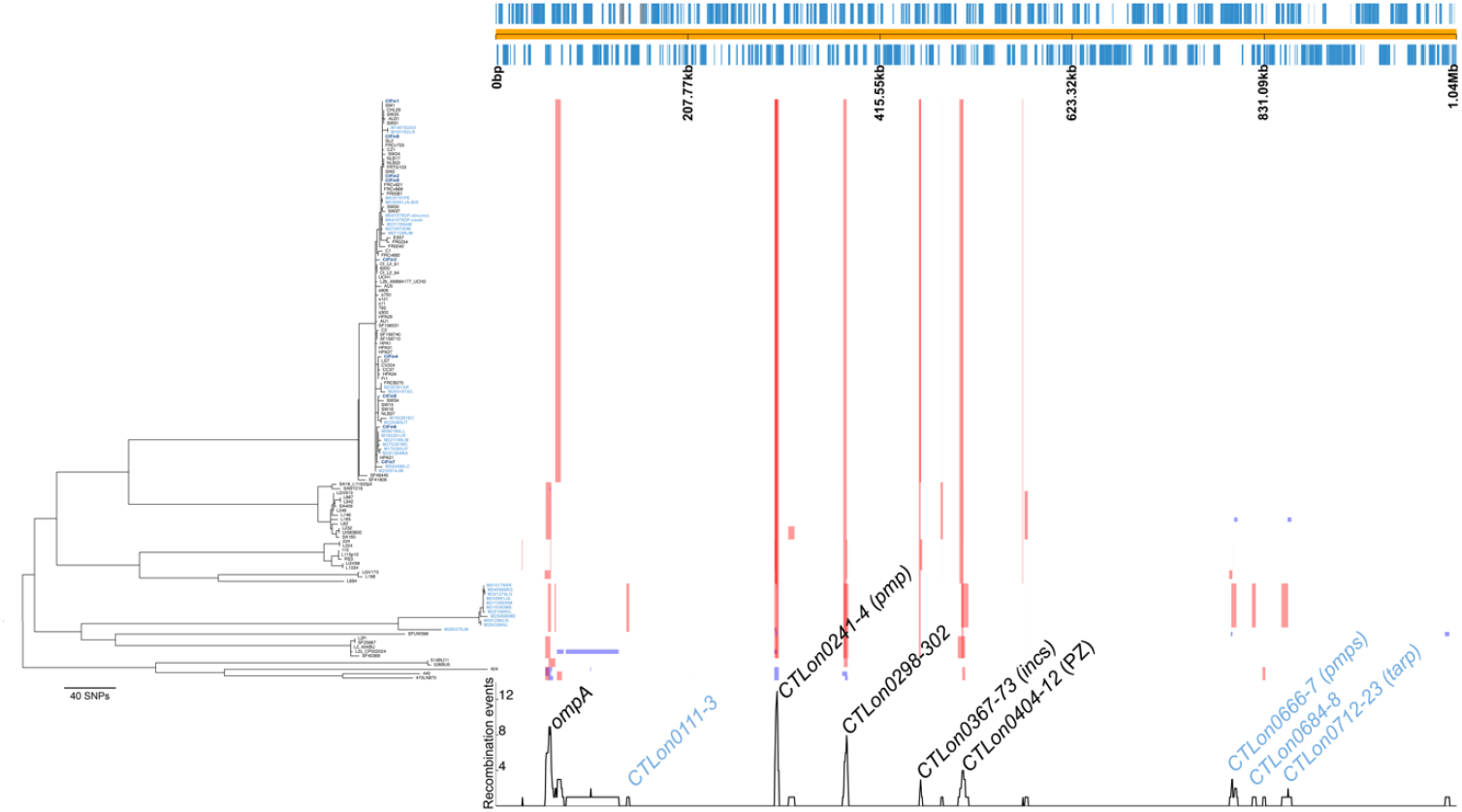
Recombination-adjusted phylogeny of LGV strains from Argentina, Finland and from previous publications showing identified recombinations. Right of the phylogeny the tracks of identified recombination are shown, with red indicating presence in multiple genomes and blue in single genomes. Above the recombination tracks are the coding sequences of strain L2b/UCH2 (AM884177) and below the tracks, peaks of recombination events. Loci affected (named from AM884177 annotation) are indicated, with those affecting in the Argentinian *ompA*-genotype L4 clade indicated in pale blue (the latter three only in the 10 strains in the tightly related clade). Newly sequenced isolates are shown in the phylogeny with labels in dark blue (Finland) and pale blue (Argentina). *pmp*=polymorphic membrane protein, *inc*=inclusion membrane protein, PZ=plasticity zone.

### Clinical relevance of *ompA*-genotype L4 lineage

Comparative analysis of the clinical data between the patients in clades L2b and L4 shows differences in the frequencies of some factors, although without statistical significance (Table 1). The presence of an anorectal inflammatory tumour was higher in patients infected by LGV from *ompA*-genotype L4 clade (50% compared to 25% in *ompA*-genotype L2b clade). The presence of anal ulcers was more frequent in patients infected with LGV from *ompA*-genotype L2b clade (55% compared to 22% in *ompA*-genotype L4 clade), and the presence of other concomitant STIs in addition to HIV was higher in LGV from *ompA*-genotype L2b clade (50% compared to 30% in *ompA*-genotype L4 clade). The numbers of cases analysed, however, are too low to draw meaningful conclusions.

**Table 1.** Clinical data of patients from Buenos Aires. Comparison of symptoms between LGV from clades *ompA*-genotype L2b and *ompA*-genotype L4. a: Rectal syndrome is characterized by rectal discharge, pushing, and tenesmus. b: Ulcers located in the anal region. c: A mass located around the anal area. d: Not verified, as rectoscopy was not performed. NG (*Neisseria gonorrhoeae*), TP (*Treponema pallidum*), HPV (Human Papillomavirus), HSV (Herpes Simplex Virus).

| Sample | LGV clade | Rectal syndrome<sup>a</sup> | Anal ulceration<sup>b</sup> | Severity of proctitis | Inguinal lymphadenopathy | Tumour<sup>c</sup> | HIV | Other STIs |
| --- | --- | --- | --- | --- | --- | --- | --- | --- |
| M010179RR | L4 | + | - | Moderate | - | - | + | - |
| M030991JA | L4 | + | - | Moderate | + | Anorectal | + | GC/HPV/TP |
| M040989RG | L4 | + | - | Severe | + | Anorectal | + | - |
| M051286CN | L4 | + | + | Moderate | - | - | - | HSV |
| M210393MB | L4 | + | - | Moderate | + | Anorectal | + | - |
| M211085WM | L4 | + | - | Not verified<sup>d</sup> | - | - | - | - |
| M250688ME | L4 | + | + | Mild | + | Anorectal | + | - |
| M251089VL | L4 | + | - | Moderate | + | - | + | - |
| M260278JB | L4 | + | - | Severe | + | Anorectal | + | TP |
| M280389NL | L4 | + | No data | Moderate | + | - | + | - |
| M030791PE | L2b | + | + | Severe | + | - | + | TP/HPV |
| M030991JA-BIS | L2b | + | - | Moderate | + | Anorectal | + | NG /TP/HPV |
| M040486LC | L2b | + | - | Moderate | - | - | + | - |
| M041079DP | L2b | - | + | Not verified<sup>d</sup> | + | No data | + | - |
| M071289JM | L2b | No data | + | Mild | - | - | + | - |
| M090190LL | L2b | + | + | Moderate | + | - | + | TP |
| M140182AO | L2b | + | - | Moderate | - | - | - | - |
| M150261JR | L2b | - | - | Moderate | - | - | + | NG/HPV |
| M150291EC | L2b | + | + | Severe | - | - | + | - |
| M161182LR | L2b | + | - | Moderate | + | - | + | - |
| M170285JP | L2b | + | - | Moderate | + | - | + | HPV/HSV |
| M200197AC | L2b | + | - | Mild | + | Anorectal | + | NG/TP/HSV |
| M210970DM | L2b | + | - | Severe | + | - | + | - |
| M220485JT | L2b | + | + | Moderate | - | - | + | - |
| M221189LM | L2b | + | - | Moderate | + | - | + | NG |
| M230381AR | L2b | + | + | Moderate | + | - | - | - |
| M230574JM | L2b | + | + | Severe | - | - | + | - |
| M231186AM | L2b | + | + | Severe | + | Anorectal | + | NG |
| M270381MC | L2b | + | + | Not verified<sup>d</sup> | + | Endoanal | + | HPV |
| M281284MA | L2b | - | + | Severe | - | Perianal | + | TP |
| p value L4/L2b |  | 0.15 | 0.11 | 0.82 | 0.55 | 0.12 | 0.65 | 0.26 |

### Urogenital T1 lineage sample causing proctitis

One sample from Argentina falls within the UG T1 lineage, as described by Hadfield *et al*. (14) (Figure 3). Sample M160595IY is located at the base of a clade of diverse genomes, from diverse countries. Sample M160595IY was identified as *ompA*-genotype D by RFLP, and from WGS is 1bp (synonymous SNP at amino acid 212) from other described D strains (Table S1). This branch is also defined by several recombination events, covering the same loci which are often affected by recombination in the T1 lineage strains (data not shown).

**Figure 3.**
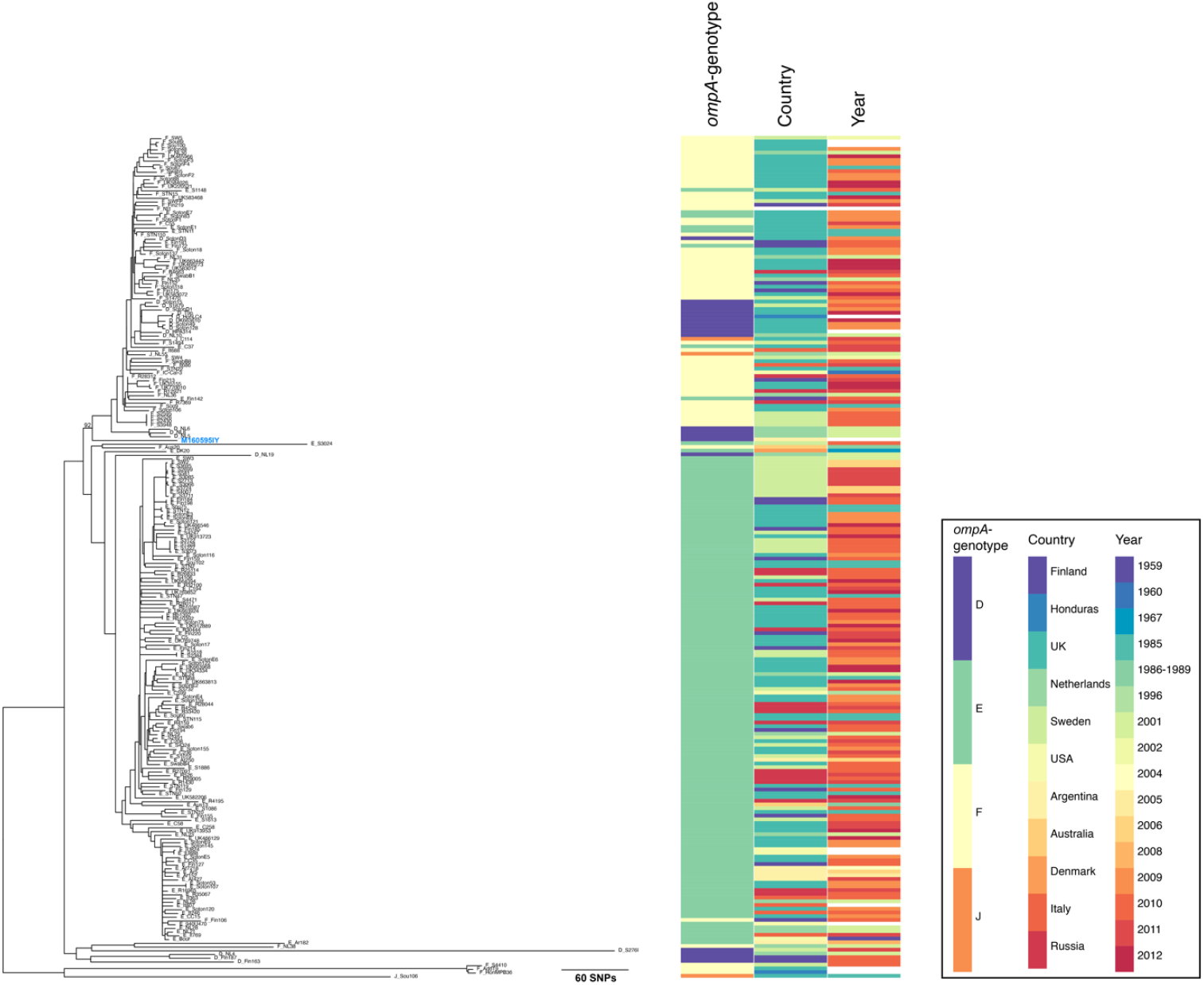
Recombination-adjusted phylogeny of UG strains belonging to lineage T1 (14) showing the location of sample M160595IY from Argentina in the context of those from previous publications. The tree is rooted according to (14). Right of the phylogeny is relevant metadata: *ompA*-genotype, country, year as described in the key to the far right. The name of the genome from this publication is pale blue in the phylogeny. The bootstrap of 92% from 1000 bootstraps is shown on the branch of interest.

Sample M160595IY was taken from a young man presenting with symptoms of mucorrhoea, proctalgia, and tenesmus, which had been developing for a month prior to the medical consultation. Diagnosis revealed mild proctitis. The patient’s sexual practices include both receptive and insertive anal sex, as well as oro-anal sex, with a reported 20 sexual partners in the previous 12 months. The patient did not present any other bacterial STI coinfections at the time of symptom onset, and responded to the antimicrobial treatment. That the genome derives from an anorectal swab from a patient with proctitis, with no evidence of a mixed infection with an LGV strain in the sample, suggests that this UG strain was the cause of mild proctitis.

Genome sequences from *ompA*-genotypes E, G and Ja isolated from rectal swabs have previously been published (21,22), and non-LGV genotypes causing proctitis in patients have been identified through *ompA* genotyping (6,23). Additionally, recombinant LGV/UG strains have been identified in the past (3,24,25); however in this case we see a genome with *ompA*-genotype D in a UG genomic backbone isolated from an anorectal swab from a male patient with proctitis. This is the first genome sequenced from a symptomatic proctitis case caused by a non-recombinant *ompA*-genotype D isolate.(6,21–23) Deeper investigation into such samples may lead to the identification of genomic regions involved in rectal tropism.

### Plasmid analysis

Plasmids from the new strains were also analysed in the context of previously published data. Among the LGV plasmids, there is very little diversity, and the plasmid is highly conserved. All the plasmids from the novel *ompA*-genotype L4 clade are located very closely in the phylogeny, with the plasmid from L1/440, L2/514BU11 and L2/526BU5, all from the USA isolated in 1968 (Figure S3A). This could indicate plasmid exchange between *ompA*-genotypes within the Americas, although further data would be required to confirm this. While the plasmids of many strains within the *ompA*-genotype L2b clade are almost identical to that in L2b/UCH2, one (from sample M230574JM) stands out as having a single, synonymous SNP in plasmid coding sequence *CDS1*.

The plasmid from the UG strain M160595IY is located in the phylogeny close to many plasmids from *ompA*-genotypes D, E, F, and J (Figure S3B). It carries three additional SNPs, representing synonymous mutations in *CDS1* and *CDS8*, and R324I in *CDS3*. This would be consistent with the genomic phylogenetic position.

**Figure S3.**
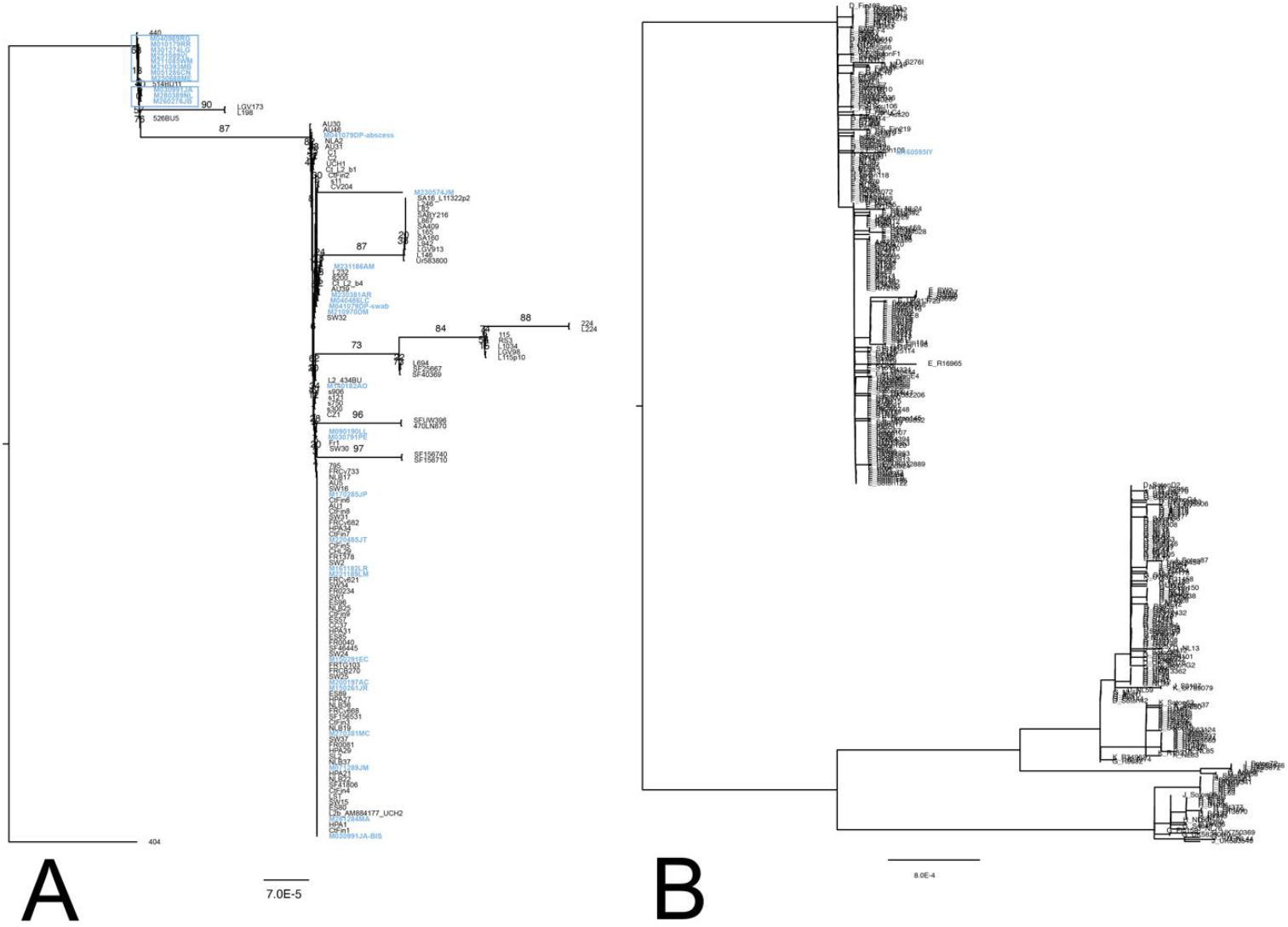
Plasmid phylogenies. **A:** Phylogeny of LGV plasmids. Rooted on L3/404. Plasmids from the *ompA*-genotype L4 clade are indicated in boxes. **B:** Phylogeny of UG plasmids. Rooted on the long branch between T1 and T2 according to UG genomes. Bootstrap values are from 1000 bootstraps.

## Conclusions

This study demonstrates the widespread global distribution of the genomic clade of *ompA*-genotype L2b, its success as the primary causative agent of LGV, and the recently evolved diversity within the clade. It confirms that in locations distant from the most studied European part of the outbreak such as Argentina, strains identified as L2 by *ompA*-genotyping also present the genetic backbone of *ompA*-genotype L2b. Our data suggests that in Buenos Aires, there have been multiple introduction events of this *ompA*-genotype L2b clade.

We also present a complete genome of a non-LGV *ompA*-genotype D sample causing proctitis, confirming their UG genomic backbone. This highlights the fact that some non-LGV strains can cause proctitis that in some cases is symptomatically similar to that produced by LGV strains.

A further key finding is the detection of a new clade causing LGV in Buenos Aires, which is vastly different from the previously described lineages, leading us to propose its designation as *ompA*-genotype L4. Future studies will further investigate the diversity within this clade and the association of the new *ompA*-genotype L4 clade with possible specific clinical manifestations.

## Author contributions

KB: Investigation, Data curation, Formal Analysis, Methodology, Visualization, Writing – original draft

FW: Investigation, Data curation, Formal Analysis, Visualization

VB: Investigation

ACE: Resources

MLGV: Resources

DLA: Resources

LLR: Resources

LSL: Resources

MHP: Resources

EHB: Resources

FI: Resources, Investigation

AE: Resources, Writing – review & editing

HMBSS: Data curation, Funding acquisition, Methodology, Project administration, Supervision, Validation, Writing – original draft

MRF: Conceptualization, Funding acquisition, Project administration, Writing – review & editing

## Ethical approval

The study on Argentinian samples was approved by an Ethics Committee (“Detección de C. trachomatis en pacientes con rectitis infecciosa: prevalencia y tipificación”, Gobierno de la Ciudad de Buenos Aires, no. 201723) and patients provided written informed consent. The Finnish samples are deidentified, and data anonymised, and therefore no specific ethical approval is required.

## Acknowledgements

We thank Daniel Gander, Valéria Pires, and Stefan Antener (all UZH) for excellent technical assistance with sequencing. We also thank Dr Osvaldo Degregorio (UBA) for his collaboration in the statistical analysis. Many thanks to Sofia Polcowñuk for designing the graphical abstract.

